# Substantial decline of phasic dopamine signaling in senescent male rats does not impact dopamine-dependent Pavlovian conditioning

**DOI:** 10.1101/2023.12.21.572806

**Authors:** Stefan G. Sandberg, Christina A. Sanford, Paul E. M. Phillips

**Affiliations:** Center for Neurobiology of Addiction, Pain & Emotion, University of Washington, Seattle, WA 98195, USA; Department of Psychiatry & Behavioral Sciences, University of Washington, Seattle, WA 98195, USA; Department of Pharmacology, University of Washington, Seattle, WA 98195, USA; Cajal Neuroscience Inc., Seattle, WA 98109, USA

## Abstract

Normal aging is associated with cognitive decline which impacts financial decision making. One of the underlying features of decision making is probability estimation, in which nucleus accumbens dopamine signaling has been implicated. Here we used fast-scan cyclic voltammetry to probe for age differences in dopamine signaling, and pharmacological manipulation to test for age differences in the dopamine dependence of Pavlovian conditioning. We found differences in phasic dopamine signaling to reward delivery, and unconditioned and conditioned stimuli, but no difference in conditioned approach between adult and senescent groups. In addition, we found that dopamine receptor antagonism with flupenthixol (225 μg/kg, i.p.) partially inhibited conditioned approach in the adult group, whereas it completely blocked conditioned approach in the senescent group. Further increase in concentration to 300 μg/kg, i.p. resulted in complete inhibition of conditioned approach behavior in both age groups. Therefore, while phasic dopamine signaling in the nucleus accumbens of senescent animals is greatly diminished in concentration, these animals maintain dopamine dependent Pavlovian conditioning.

## Introduction

The pathological process of aging is well associated with the neurodegenerative conditions resulting in Alzheimer’s (AD) and Parkinson’s disease (PD) among others. However, the normal process of aging, known as senescence, is associated with non-pathological neuronal loss and cognitive decline as well (Juarez et al., 2019). While the symptoms of AD and PD are debilitating, senescence by itself present challenges to the aging population, for example in decision making. Control of decision making can be rationalized into multiple systems with different forms of computations, labeled as model based and model free (Dayan and Daw, 2008). The model-based system(s) is a computationally demanding and flexible system, thought to rely on cognitive representations of stimuli in an environment predicting an appetitive outcome. Conversely, a model-free system has a low computational demand and utilizes cached based values to guide an animal towards an appetitive outcome. In the model-free system, cache values are assigned to appetitive cues, values that in turn are updated through experience via mechanisms thought to follow temporal difference learning algorithms (Sutton and Barto, 1987; Schultz et al., 1997) based upon formal learning theories (Rescorla Wagner, 1972). Previous work has identified dopamine signaling as a potential neural substrate for the model-free system and, in particular, phasic dopamine signaling (Flagel, Clark et al, 2011), has been implicated in aspects of associative learning and decision making (Gan et al., 2010; Saunders et al., 2018; Sharpe et al., 2020). Moreover, other studies have shown biochemical (Morgan et al., 1987; Moretti et al., 1987; Aria and Kinemuchi, 1988; Gozlan et al., 1990; Suzuki et al., 2001), neurochemical (Yurek et al., 1998;Friedemann and Gerhardt, 1992) evidence indicating a decline in dopamine signaling associated with senescence. These two observations (dopamine’s involvement in decision making and associative learning, and age related degeneration of dopamine signaling) could potentially explain the cognitive decline due to senescence. Specifically, if degenerated dopamine signaling is manifested as reduced phasic dopamine signaling, we would anticipate a shift away from the dopamine dependent model-free decision making in favor of higher dependence upon model-based decision-making processes. Conversely, in the absence of such a switch one would anticipate behavioral deficits in decision making and/or associative learning. To this point, biochemical, neurochemical, and behavioral changes may be to culprit to altered risk-based decision making that has been shown to be behaviorally different across the lifespan in humans. When probabilities of outcomes are known, older adults are less risk taking than young adults (Deakin et al., 2004). In addition, using data on real financial portfolios of individual investors, Korniotis & Kumar (2011) found that older adults do not reduce their exposure to the stock market as a whole, but tend to choose less risky stocks. Data from the Survey of Consumer Finances, however, indicate that individuals over the age of 65 are less willing to take high risks to secure large financial returns, also when faced with hypothetical situations. For people younger than 65 there is no clear correlation between risk tolerance and age (Poterba, 2001). Importantly, older and younger adults seem to employ different strategies when making risky choices, especially when faced with ambiguity (e.g., Iowa Gambling Task) even when their net performance on the task is similar. Younger adults exhibit negativity bias (i.e., are influenced more by past losses) but recall past outcomes better than older adults (Wood et al., 2005). Older adults are considered to have more difficulty in detecting the complexity of risky choices and, consequently, often choose the risk-less, simple option. This could be due to a lack of dynamic range when describing a probabilistic outcome, that is to say having a decreased resolution of encoding finely separated probabilistic outcomes and in turn lead to maladaptive associative representations of risky outcomes. While risk based decision making in the financial market is a complex behavior, there are data suggesting a connection between dopamine phasic signaling and the development of experience dependent estimation of probabilities and risk, in accordance with model-free mechanisms (Nasrallah et al., 2011; Clark et al., 2012). Given the biochemical, neurochemical and pharmacological evidence of dopamine signaling loss associated with senescence, and observed human behavioral changes associated with aging we wanted to investigate whether changes in phasic dopamine signaling due to aging was present, and for possible behavioral deficits associated with aging in a rodent model by utilizing fast-scan cyclic voltammetry, and pharmacology during Pavlovian conditioning.

## Methods

### Animals and Surgery

Animals used in all experiments were male Fisher F-344 (Charles River Laboratories), where adult group were 6-8 months in age and senescent group was 24-26 months in age. Rats were pair-housed upon arrival, maintained on a 12-hr light/dark cycle, and given ad libitum access to water and lab chow. In order to start behavioral testing and voltammetric measurements at 6 and 24 months, respectively, we performed surgeries at month 5 and 23 months (300-400g). Surgeries have been previously described (Clark, Sandberg et al., 2010), but briefly animals were induced with isoflurane at 5% animals were then placed in stereotaxic apparatus where deep anesthesia was maintained at 1-3% isoflurane, scalp was swabbed with 70% isopropanol and 10% providone pads. Subsequently, scalp was injected with bupivacaine and lidocaine solution. Scalp was then removed and skull exposed and holes drilled for anchor screws, Ag/AgCl reference, AP:-2.0; ML -3.0, and working electrodes, AP: +1.3; ML: +1.3; DV: -7.0 (all measurements in mm). In order to protect exposed skull and to secure electrodes they were covered in acrylic dental cement (Lang Dental). Post recovery, 1 week, animals were placed on food deprivation where animals were kept at 90% body weight (assuming 1.5% bodyweight increase per week) throughout behavioral testing.

### Pavlovian Conditioning

Pavlovian conditioning consisted of 10 sessions with 25 trials per session. Each trial consisted of a cue delivery (tray light as conditioned stimulus, CS) lasting 8 s, followed by pellet delivery (unconditioned stimulus, US, Bio Serv). The inter trial interval was 60±30s at 10s intervals. Conditioning was performed in extra tall 5 panel operant boxes from Med Associates. Boxes were equipped with food trays and cue lights above food trays. Before initiating Pavlovian conditioning, animals were pre exposed to food pellets in order to eliminate neophobic behaviors during conditioning. Our metric of conditioned approach behavior was defined as head entries into food tray and quantified with photo beam breaks. Difference score was defined as rate (s^-1^) of head entry during CS – rate (s^-1^) of head entry during inter-trial interval. We also measured reaction time to head entry after CS onset.

### Pharmacological interrogation of dopamine dependence on Pavlovian conditioned approach behavior

Here we used flupenthixol (FLU, a mixed D1/D2 receptor antagonist) to test for dopamine dependence of conditioned approach behavior in a similar manner as previously described (Flagel, Clark et al., 2010). Briefly, rats were given an injection either saline (SAL,i.p.; 0.9% NaCl) or 225□μg□kg^−1^ of flupenthixol one hour before Pavlovian conditioning sessions 1–6. This dose of drug was chosen based on a previous study, where they demonstrated that this dose was not resulting in any nonspecific inhibitory effects on motor activity (Flagel, Clark et al., 2010). In order to test for dopamine dependence of conditioned response, both groups received an injection of saline before session 7. The metric used for quantifying conditioned response was head entries into food tray, quantified as difference score (rate of head entry during CS presentation – rate of head entry during inter CS interval.

### Voltammetry recordings

During all experimental sessions, chronically implanted microsensors were connected to a head-mounted operational amplifier for current-to-voltage transduction allowing for dopamine detection utilizing FSCV. Real time measurements of dopamine levels were performed at a sampling frequency of 10□Hz. Measurements of dopamine are done by monitoring oxidative and reductive currents caused by the application of a voltage triangle waveform (a scan), when dopamine is present at the surface of the electrode during a voltammetric scan, it is oxidized during the anodic sweep to form dopamine-*o*-quinone (peak reaction at approximately +0.7□V), which is reduced back to dopamine in the cathodic sweep (peak reaction at approximately −0.3□V). The measured oxidative and/or reductive current is directly proportional to the number of molecules, thus yielding quantitative information. In addition, maximum oxidative and reductive currents are dependent on applied potential, and the rate at which it is applied or scanned. This in turn provides a chemical signature that is characteristic of the analyte, allowing resolution of dopamine from other electroactive substances. For quantification of changes in dopamine concentration over time, the peak oxidative current were plotted for successive voltammetric scans. Waveform generation, data acquisition and analysis were carried out on a PC-based system using two PCI multifunction data acquisition cards (NI PCI-6052E and NI PCI-6711) and software written in LabVIEW (National Instruments).

### Statistical analysis of voltammetry data

Data analysis of voltammetric data was carried out using software written in LabVIEW (National Instruments) and low-pass filtered at 2 kHz. Dopamine was extracted from the voltammetric signal with a calibration method known as chemometric analysis (Rodeberg et al., 2017) using a standard training set based upon stimulated dopamine release of freely moving rats detected at chronically implanted electrodes. In this calibration procedure dopamine concentration was estimated on the basis of the average post-implantation sensitivity of electrodes. All data were smoothed with a 5-point within-trial running average. Peak dopamine values in response to the US and CS were obtained by taking the largest value in the 3-s period after stimulus presentation. Peak values were then compared using two separate mixed models ANOVAs (one for CS and one for US) with session as the repeated measure and age (adult vs. senescent) as the between-group measure. All post-hoc comparisons were made with the Bonferroni correction for multiple tests. All statistical analyses were carried out using Prism (GraphPad Software).

### Histological verification of recording site

After behavioral testing was over, animals were anesthetized with ketamine (100□mg□kg^−1^, i.p.) and xylazine (20□mg□kg^−1^, i.p.) and then transcardially perfused with saline followed by 4% paraformaldehyde. Brains were removed and post-fixed in paraformaldehyde for 24□h and then frozen in -80 ºC, sliced on a cryostat (50-μm coronal sections, -22 --25□ºC) and stained with cresyl violet to aid in visualization of anatomical structures. Electrode placements can be viewed in Fig. 1.

**Figure 1.**
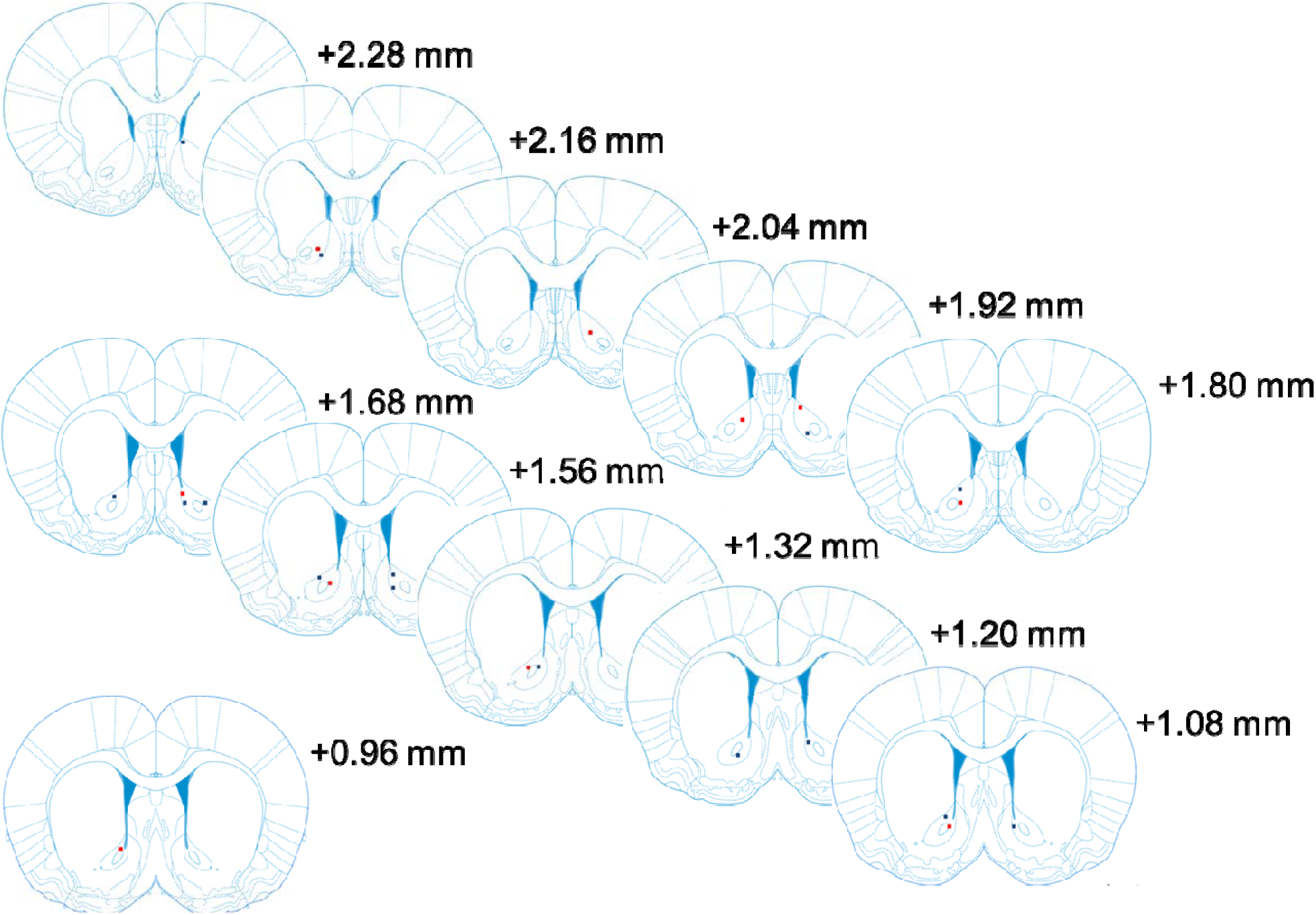
Recording locations from the nucleus accumbens core. Anterior posterior coordinates are next to each brain section, given in millimeter. Adapted from Paxinos G, Watson C (2005) Rat Brain in Stereotaxic Coordinates (Elsevier Academic, Burlingame, MA) 5th Ed.

## Results

### Nucleus accumbens core dopamine release to unexpected pellets and Pavlovian stimuli is decreased in senescent population

In order to test whether biochemical markers indicating decreased dopamine biosynthesis resulted in decreased dopamine signaling we implanted adults (6-8 months) and senescent (24-26 months) Fisher 344 rats with biocompatible carbon fiber microsensors (Clark and Sandberg et al, 2010). These sensors enabled longitudinal dopamine measurements across the time span of Pavlovian conditioning learning. Before each session animals were tested to see if dopamine signals were sensitive to reward magnitude and if these signals differed between age groups. As can be seen in the individual representative traces in Fig. 2*a* and *b* dopamine signals appeared to reflect reward magnitude along with greater increase to 1 and 4 pellets in the adult compared to the senescent animal. Indeed, when summarized across all animals and statistical comparison between the age groups were done, there were significant increases in signal when delivering 1 versus 4 pellets (F_1,22_=7.167, p=0.0138), in addition there was a significant difference between age groups (F_1,22_ =14.47, p=0.001). While delivery of unexpected pellets is of behavioral significance, we were also interested in whether observed age group differences were present in behaviorally evoked signals associated with Pavlovian conditioning, i.e. CS and US. In this Pavlovian task the CS (cue light in the food hopper) was on for 8s and terminated upon the delivery of one food pellet (US). Adult and senescent groups are summarized in pseudo 3D plots, Fig. 3*a* and 3*b*, respectively. As can be seen from these figures, the adult group display bigger dopamine signals overall with visible dynamic changes, i.e. signals being greater to the US in the early trials (up to <25 trials), and US signals diminishing and CS signals increasing towards later trials (>50-75 trials). In contrast, signals to CS and US in senescent group are smaller and lack dynamic changes. These qualitative and quantitative changes were further summarized by extracting individual voltammetry recordings from each Pavlovian trial with chemometrics, and peak dopamine signals from CS and US were calculated as the difference between pre-stimulus baseline and 1 s averaged at peak signal post stimulus, Fig. 3*c*. Previous observations from pseudo 3D plots were statistically confirmed by comparing session averages between age groups for both CS and US. As can be seen from the group averaged FSCV traces, the adult group CS signals increased with increasing experience (main effect of session, F_(2.774, 44.38)_ = 4.131, p=0.0133), and were significantly different between the two age groups (main effect of age, F_(1, 16)_ = 18.92, p=0.0005), albeit in a different way between the two age groups (main effect of interaction, F_(5, 80)_ = 2.786, p=0.0227), Fig. 3*b*. In addition, these same statistical effects were present in the US signals as well, Fig. 3*c*. Here, though, signals decreased with experience (main effect of session, F_(1.751, 28.02)_ = 9.971, p=0.0008), age (main effect of age, F_(1, 16)_ = 18.59, p=0.0005), with a different pattern of change in US signals between the two age groups (main interaction effect, F_(5, 80)_ = 10.71, p<0.0001).

**Figure 2.**
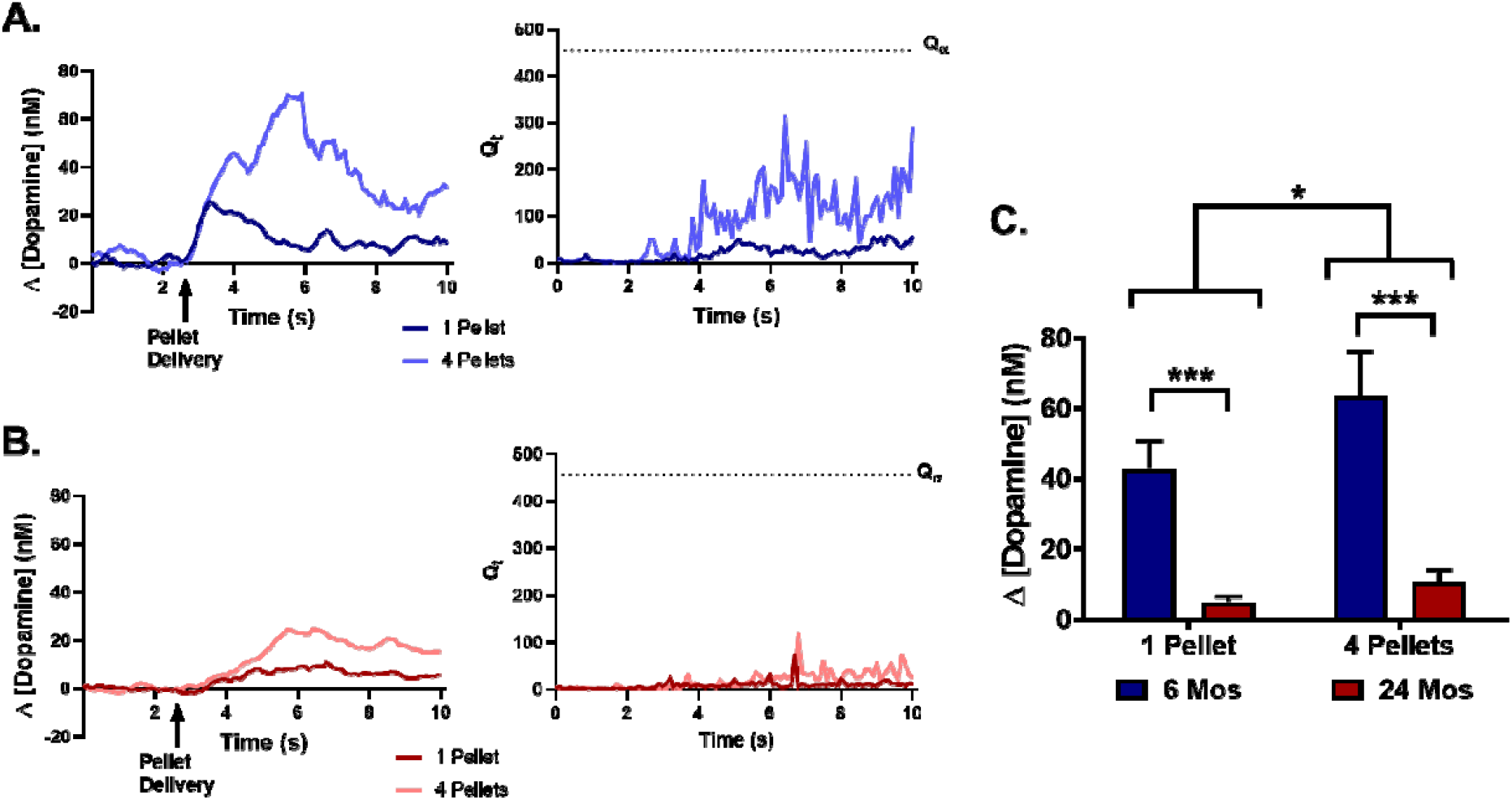
Example traces from unexpected pellet delivery of one and four pellets from individual animals of both age groups **A**. 6 months, adult and **B**. 24 months, senescent. Left panels are the dopamine signals as extracted with chemometrics. Right panels are the corresponding timeseries of residuals (Q_t_) from each individual trace. As can be seen at no point does the Q_t_ cross the critical threshold Q_α_, indicating that statistical model used to extract dopamine signal is not violated. **C**. Summarized data from both age groups and number of pellets delivered. There was a main effect of age (F_(1, 22)_ = 14.47, p=0.0010) and pellet number (F_(1, 22)_ = 7.167, p=0.0138) along with a significant interaction effect (F_(1, 22)_ = 2.272 P=0.1459).

**Figure 3.**
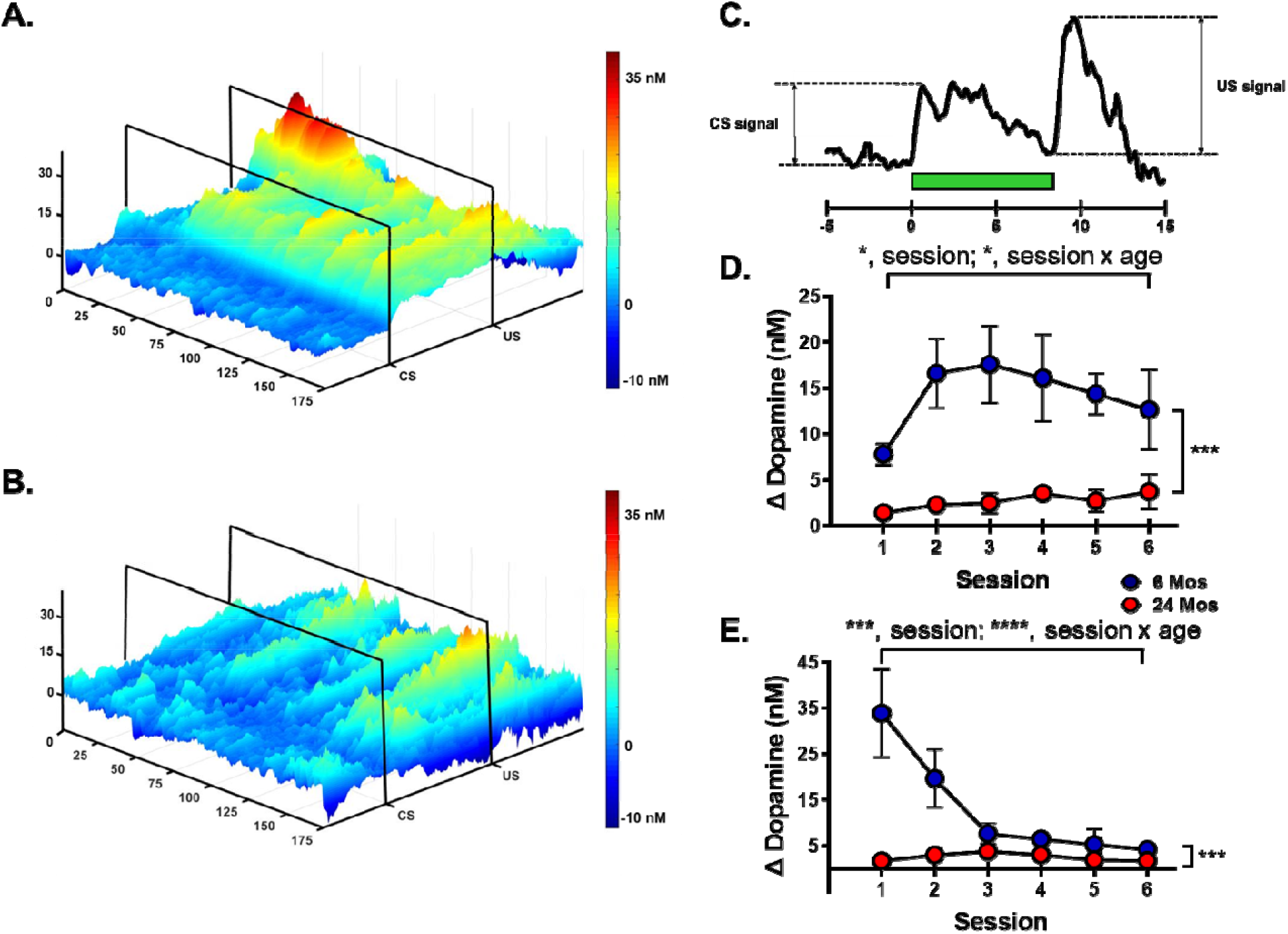
Adults have greater behaviorally evoked dopamine signals and display learning related changes in dopamine release dynamics as compared to senescent group. **A**. Summary of voltammetry data from Pavlovian conditioning between adult (n=7), and senescent (n=11). Averaged data of 175 trials with 2D 5-point smooth from adult group, and **B**. senescent group. **C**. Example trace illustrating how CS and US signals have been extracted in subsequent figures. **D**. Summary of CS signals and over 6 sessions from adult and senescent groups. There were significant main effects of age (F_(1, 16)_ = 18.92, p=0.0005), session (F_(2.774, 44.38)_ = 4.131, p=0.0133), and interaction of age and session (F_(5, 80)_ = 2.786, p=0.0227). **E**. Summary of US signals and over 6 sessions from adult and senescent groups. There were significant main effects of age (F_(1, 16)_ = 18.59, p=0.0005), session (F_(1.751, 28.02)_ = 9.971, p=0.0008), and interaction of age and session (F_(5, 80)_ = 10.71, p<0.0001).

### No behavioral difference between age groups, despite differences in dopamine signals

Because we have now an age difference in dopamine not only to unexpected pellets, but also to behaviorally evoked signals throughout Pavlovian conditioning we wanted to analyze for potential age associated differences in conditioned approach behavior. Here statistical analysis indicated that there was a significant effect of session (F_3.676, 58.81_=16.81, p<0.0001) suggesting an increase in conditioned approach behavior, consistent with learning, Fig. 4*a*. However, no significant difference was found between age groups (F_1,16_=0.1673, p=0.6879), this despite the fact that dopamine signaling was different in magnitude and learning associated changes over time between the two age groups. Next, we analyzed reaction time to first head entry upon CS presentation. Similar to conditioned approach behavior, we found that there was a significant effect of session (F_3.324, 53.18_=5.319, p=0.0021), and no difference between age groups (F_1,16_=1.12, p=0.3057), Fig. 4*b*. Given these surprising results, i.e. a difference in dopamine signaling to CS and US’s and no difference in behavior, we decided to test for dopamine dependence of the Pavlovian conditioning between the two age groups.

**Figure 4.**
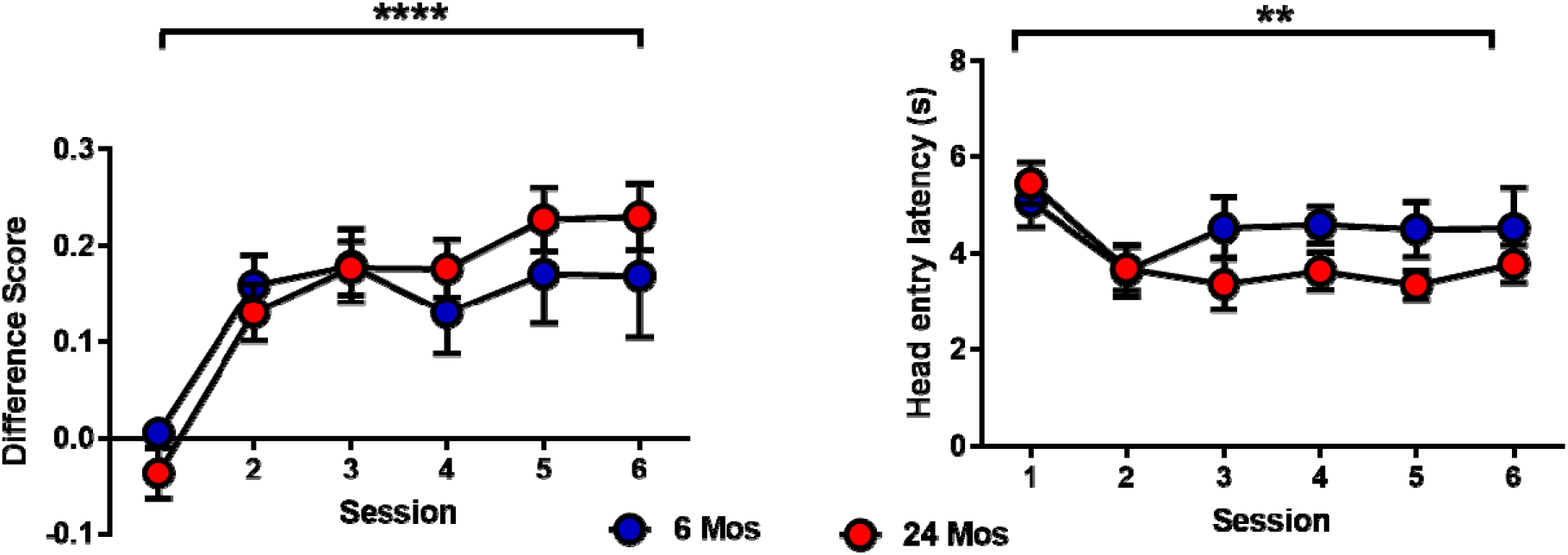
No difference in conditioned response between adult and senescent groups. Behavioral results from Pavlovian conditioning compared between adult (n=7), and senescent (n=11). **Left panel**, conditioned approach as measured by difference score (DS, rate of head entry during CS presentation – rate of head entry during inter CS interval). **Right panel**, latency to head entry (LHE) into food hopper upon CS presentation. No significant difference between age was observed in DS metric (F_(1, 16)_ = 0.1673, p=0.6879) and HEL (F_(1, 16)_ = 1.120, p=0.3057), whereas there was a significant effect of session in DS metric (F_(3.676, 58.81)_=16.81, p<0.0001), and HEL (F_(3.324, 53.18)_ = 5.319, p=0.0021). No significant interaction was found for either behavioral measurements either, DS (F_(5, 80)_ = 1.314, p=0.2666), or HEL (F_(5, 80)_ = 1.806, p=0.1211).

### Differential sensitivity to dopamine antagonism of conditioned response between age groups

In order to test for dopamine dependence of Pavlovian conditioning, we employed the approach previously described by Flagel, Clark et al. (2012), but with a total of 7 Pavlovian sessions as opposed to 8. Each age group received either intraperitoneal injections of SAL or FLU (225 μg/kg) before each session during the conditioning phase (session 1-6) and for the 7^th^ session they all received SAL, in this way we can test for dopamine’s involvement in the learning of conditioned approach behavior. Based on the difference in dopamine signals with no difference in approach behavior between the two age groups, we predicted that the senescent group would display dopamine independent learning and adult group would display dopamine dependent learning. This experiment was analyzed with a three-way ANOVA, since we had a 2x2x6 design (age x drug x sessions), Fig. 5*a*. SAL treated animals of both age groups displayed learning as the dependent variable was significantly dependent on session, F_(2.831, 79.26)_ = 45.96, p<0.0001. There was also a significant main effect of FLU treatment, F_(1, 28)_ = 20.36, p=0.0001, whereas no main effect of age was observed, F_(1, 28)_ = 0.07198, p=0.7904. More interesting patterns emerged when we looked at the interaction of main effects. In the two-way interactions we found a significant main effect of session and drug (F_(5, 140)_ = 9.599, p<0.0001), this was interpreted as the FLU effect being dependent on session, meaning that there is partial blockade of conditioned approach in FLU treated animals as compared to conditioned approach observed in SAL treated animals. The three-way interaction effect between age, drug, and session was also significant, F_(5, 140)_ = 5.909, p<0.0001, which suggested that not only did the approach behavior develop differently over time, depending on whether FLU was present or not, but also that the effect of FLU was dependent on age group as well. To confirm this, we employed a different statistical analysis, linear regression, where the two FLU treated age groups showed a separation, as indicated by a significant increase in conditioned approach in the adult group FLU treated group and not in the senescent group, r^2^=0.4273, p<0.0001, and r^2^=0.0152, p=0.3744, respectively. When animals were then tested on the 7^th^ session with SAL across all treatment groups, we found that there was a main effect of FLU history (i.e., whether the animals had received SAL or FLU throughout the 6 preceding conditioning sessions), F_(1, 28)_ = 13.58, p=0.0010, Fig. 5*b*. Post hoc analysis with Bonferroni correction for multiple comparisons revealed that this main effect was driven by a significant difference between SAL and FLU in the senescent group, t_(14)_=3.428, p=0.0038, as no significant difference was observed in the adult group. This result was surprising as it suggested that the adult group was dopamine independent in this Pavlovian paradigm. Alternatively this result could be due to there being a difference in dopamine antagonism between the age groups because of the different levels of dopamine signal present during conditioning in both age groups. In order to test whether the adult group was dopamine independent or the alternative explanation was present we increased the FLU dose (300 μg/kg, i.p.) to a new cohort of animals and repeated the experiment, Fig. 6*a*. At the 300 μg/kg dose we found a significant two-way interaction between session and drug, F_(5, 85)_ = 18.40, p<0.0001, again indicating that presence of FLU interacted with the animals’ ability to demonstrate learned changes in conditioned approach behavior across sessions. More intriguingly, there was no significant three-way interaction, indicating that there was no age-related effect on the ability of FLU to alter session dependent changes in conditioned approach behavior. Indeed, when SAL test on session 7 was performed and statistically analyzed we again found a main effect of FLU history, F_(1, 18)_ = 21.82, p=0.0002, Fig. 6*b*. Moreover, post-hoc analysis now revealed that there was a significant decrease in conditioned approach in both age groups, t_(10)_ = 3.313, p=0.0077 (adult) and t_(8)_ = 3.304, p=0.0079 (senescent). The second metric for conditioned approach, head entry latency, showed a similar pattern in the 225 μg/kg, i.p. experiment, where we saw a trend towards partial blockage of learning in adult group versus complete blockage in senescent group during conditioning phase, Fig. 7*a*. While the three-way ANOVA did not generate a three-way interaction suggestive of partial blockade of learning, the two-way ANOVA performed on the test day of the 7^th^ session did result in a main effect of drug history, F_(1, 28)_ = 5.820, p=0.0226. Post hoc analysis revealed that this main effect was driven by a significant difference between SAL and FLU treated animals in the senescent group, adult: t(14)=0.5791, p>0.9999, and senescent: t(14)=2.824, p=0.0173, Fig. 7*b*. This result further supported the idea of dopamine independence or incomplete dopamine antagonism of head entry latency in the adult group. The results become a bit more confusing when considering the 300 μg/kg, i.p. experiment. Here, the three-way ANOVA demonstrated a clear blockage of learning throughout the conditioning phase (F_(5, 85)_ = 3.132, p=0.0121, session and drug interaction), whereas the two-way ANOVA performed on the 7^th^ session test day did not demonstrate any main effect of drug history (SAL vs. FLU session 1-6), Fig. 8*a* and *b*.

**Figure 5.**
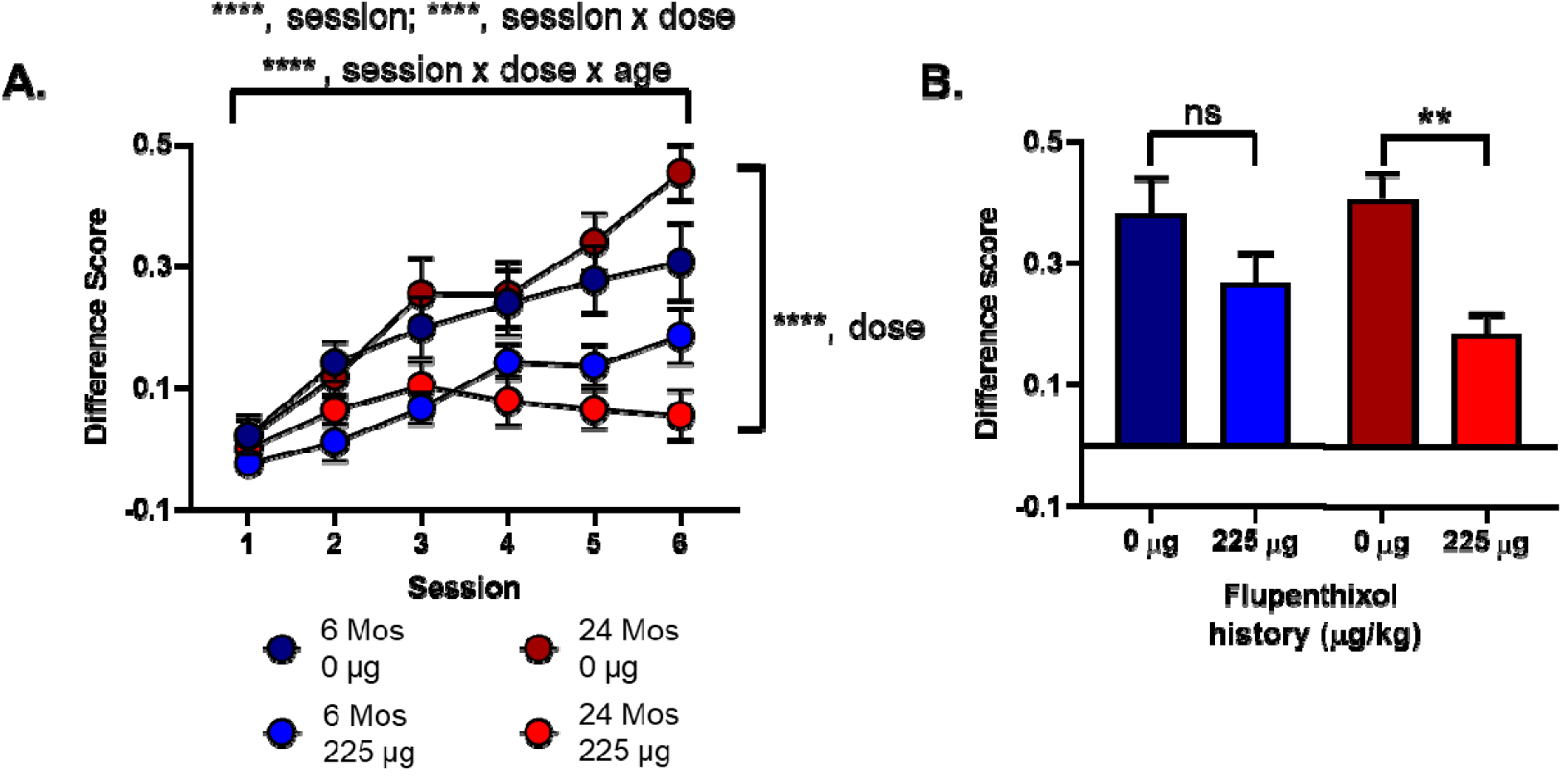
Conditioned response, as measured by difference score, was partially blocked by dopamine antagonism (Flupenthixol (FLU) 225 μg/kg, i.p.) in adult group, whereas senescent group showed a stronger block of conditioned response. **A**. 3-way comparison of conditioned approach as measured by difference score across 6 session with 225 μg/kg of FLU injected daily, 60 minutes prior to start of session. There were significant main effects of session (F_(2.831, 79.26)_ = 45.96, p<0.0001), dose (F_(1, 28)_ = 20.36, p=0.0001), session x dose interaction (F_(5, 140)_ = 9.599, p<0.0001), and session x dose x age interaction (F_(5, 140)_ = 5.909, p<0.0001). **B**. 2-way comparison of conditioned approach as measured by difference score of saline injection only (dopamine antagonist-free) session 7. There was a main effect of drug history, i.e. saline or FLU during six preceding sessions, (F_(1, 28)_ = 13.58, p=0.0010). Post hoc analysis with Bonferroni correction for multiple comparisons revealed a non-significant difference between saline and FLU treated animals in the adult group (t_(14)_=1.776, p=0.1731), and a significant difference in the senescent group (t_(14)_=3.428, p=0.0038).

**Figure 6.**
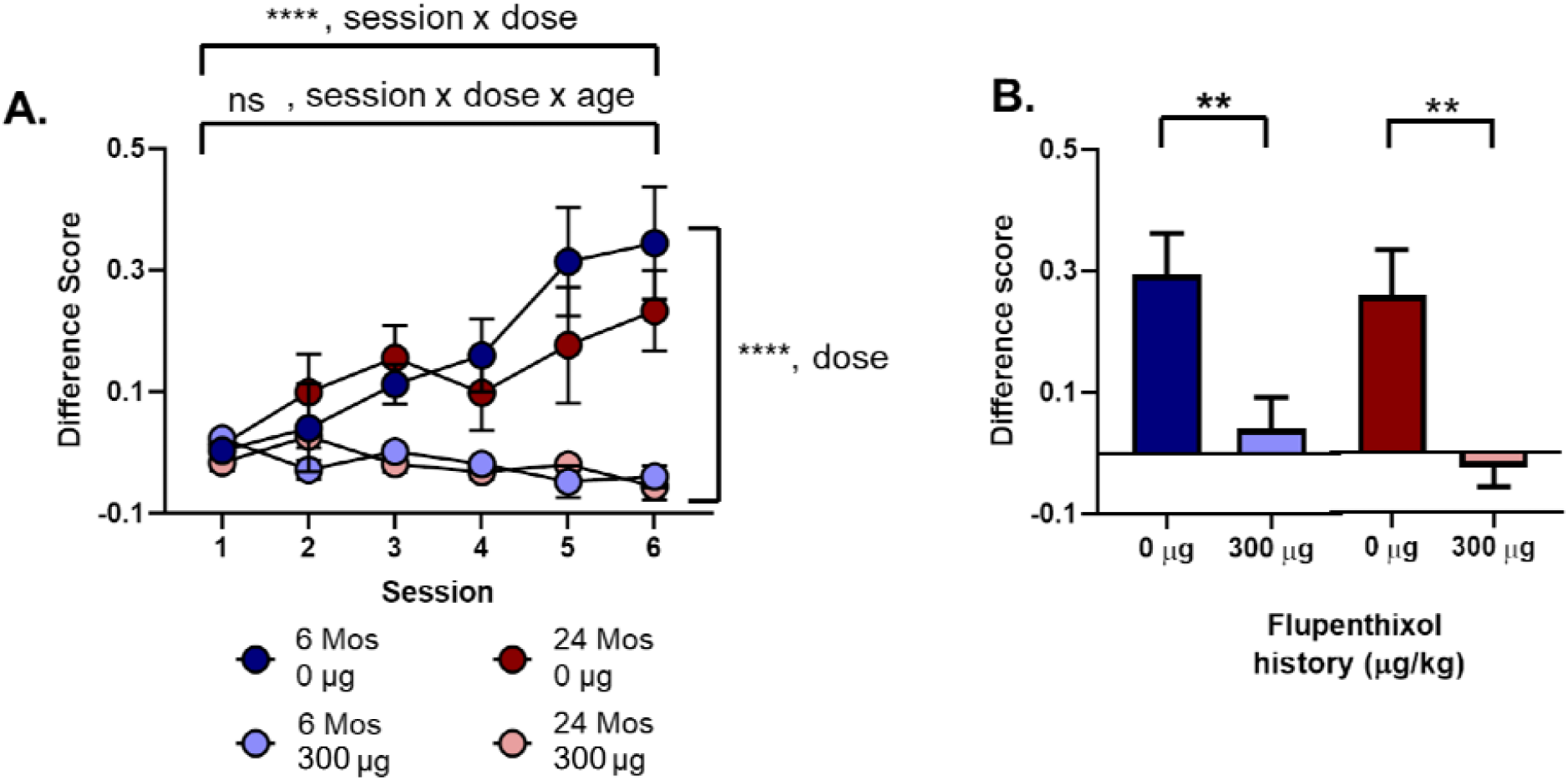
Conditioned response, as measured by difference score, was blocked by dopamine antagonism (Flupenthixol (FLU) 300 μg/kg, i.p.) in adult group and senescent group. **A**. 3-way comparison of conditioned approach as measured by difference score across 6 session with 300 μg/kg of FLU injected daily, 60 minutes prior to start of session. There were significant main effects of session (F_(2.754, 46.82)_ = 8.899, p=0.0001), dose (F_(1, 17)_ = 26.29, p<0.0001), session x dose interaction (F_(5, 85)_ = 18.40, p<0.0001). However no session x dose x age interaction was observed. **B**. 2-way comparison of conditioned approach as measured by difference score of saline injection only (dopamine antagonist-free) session 7. There was a main effect of drug history, i.e. saline or FLU during six preceding sessions, (F_(1, 18)_ = 21.82, p=0.0002). Post hoc analysis with Bonferroni correction for multiple comparisons revealed a significant difference between saline and FLU treated animals in the adult group (t_(10)_=3.313, p=0.0077), and senescent group (t_(8)_=3.304, p=0.0079).

**Figure 7.**
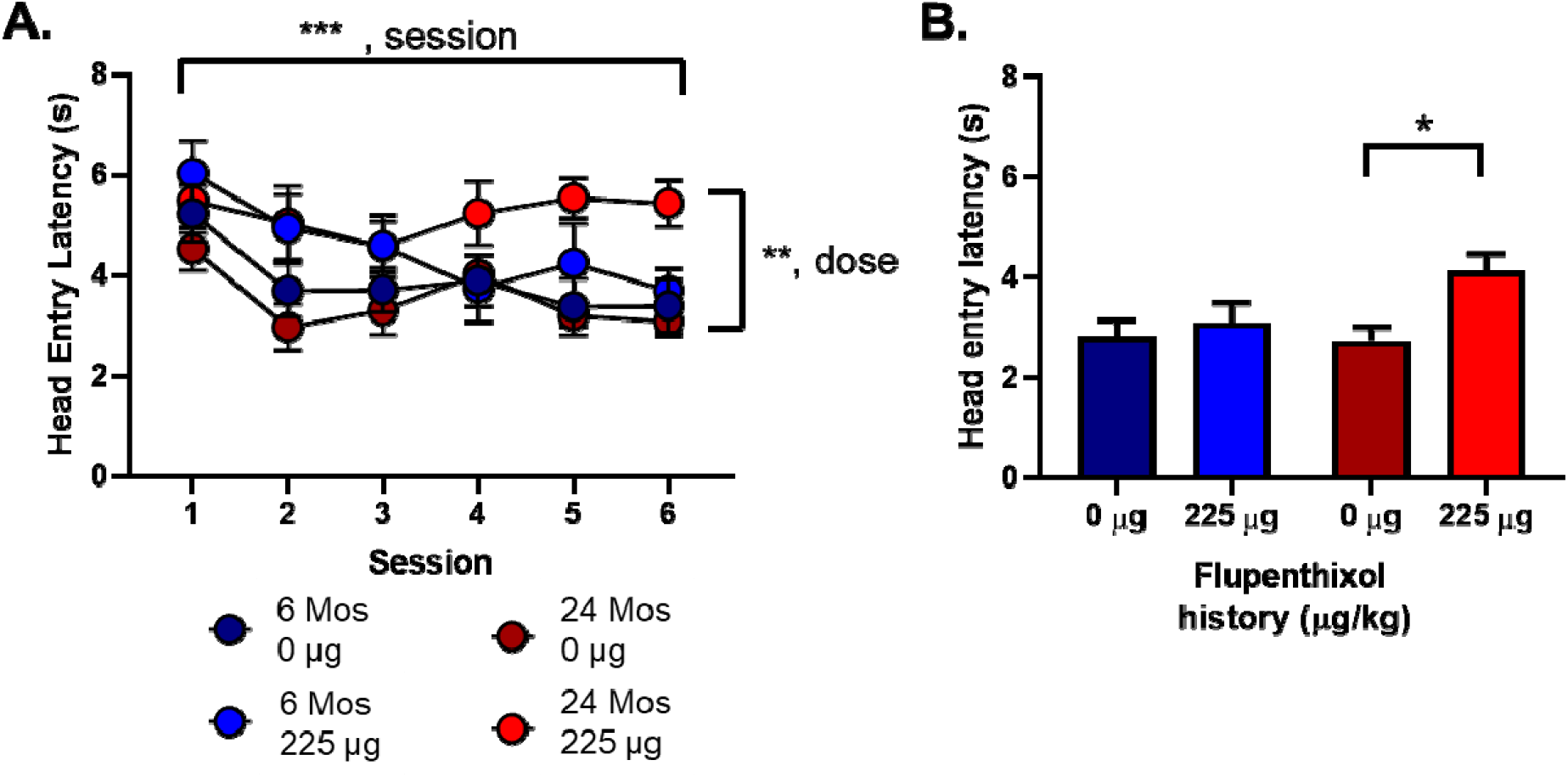
Conditioned response, as measured by latency to head entry into food hopper, was not blocked by dopamine antagonism (Flupenthixol (FLU) 225 μg/kg, i.p.) in adult group, whereas senescent group showed a blocked conditioned response. **A**. 3-way comparison of conditioned approach as measured by head entry across 6 session with 225 μg/kg of FLU injected daily, 60 minutes prior to start of session. There were significant main effects of session (F_(3.375, 94.51)_ = 6.441, p=0.0003), dose (F_(1, 28)_ = 8.225, p=0.0078), a trend toward session x dose interaction (F_(5, 140)_ = 2.262, p=0.0516), however no session x dose x age interaction (F_(5, 140)_ = 0.7654, p=0.5762). **B**. 2-way comparison of conditioned approach as measured by latency to head entry into food hopper of saline injection only (dopamine antagonist-free) session 7. There was a main effect of drug history, i.e. saline or FLU during six preceding sessions, (F_(1, 28)_ = 5.820, p=0.0226). Post hoc analysis with Bonferroni correction for multiple comparisons revealed a non-significant difference between saline and FLU treated animals in the adult group (t_(14)_=0.5791, p>0.9999), and a significant difference in the senescent group (t_(14)_=2.824, p=0.0173).

**Figure 8.**
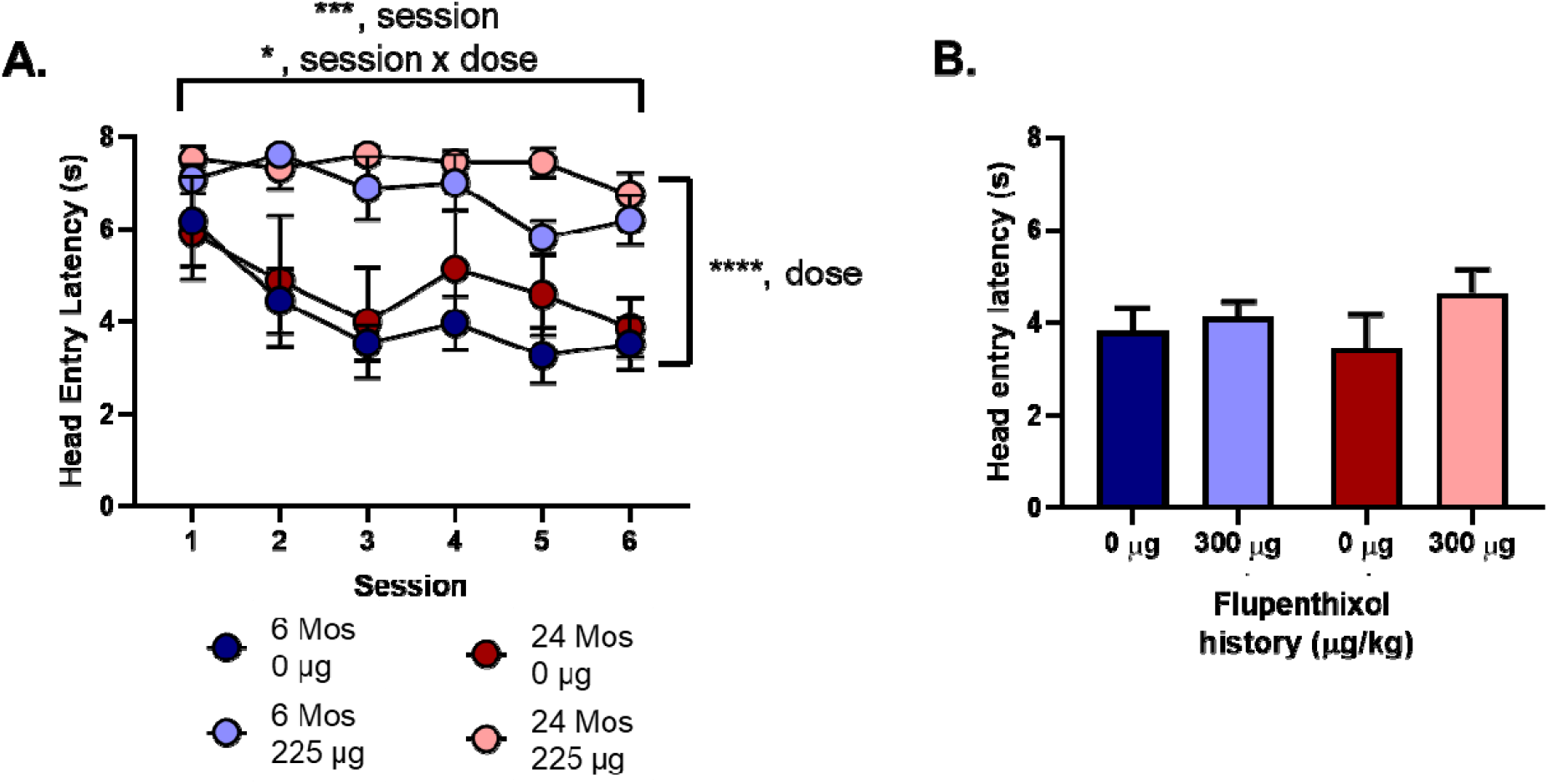
Conditioned response, as measured by latency to head entry into food hopper, was blocked by dopamine antagonism (Flupenthixol (FLU) 300 μg/kg, i.p.) in both groups during conditioning phase. **A**. 3-way comparison of conditioned approach as measured by difference score across 6 session with 300 μg/kg of FLU injected daily, 60 minutes prior to start of session. There were significant main effects of session (F_(3.011, 51.19)_ = 7.944, p=0.0002), dose (F_(1, 17)_ = 28.5, p<0.0001), session x dose interaction (F_(5, 85)_ = 3.132, p=0.0121), however no session x dose x age interaction (F_(5, 85)_ = 0.5130, p=0.7657). **B**. 2-way comparison of conditioned approach as measured by latency to head entry into food hopper of saline injection only (dopamine antagonist-free) session 7. There was no main effect of drug history, i.e. saline or FLU during six preceding sessions.

## Discussion

Here we utilized FSCV and pharmacology to investigate for the presence of an age associated decrease in behaviorally evoked phasic signaling and any accompanying behavioral deficit. Similar to previous neurochemical studies (Friedemann and Gerhardt, 1992; Yager et al.1998), we did observe a significant decrease in phasic dopamine signaling elicited by unexpected reward delivery and to cues predicting rewards delivery all within a Pavlovian conditioning context. To our surprise, this clear reduction in phasic signaling did not result in a behavioral deficit as measured by conditioned approach behavior. One possible explanation for this lack of behavioral deficit could be that loss of dopamine signaling could have led to a switch in dopamine dependence of the Pavlovian conditioning. In the study of Flagel, Clark et al. (2012), animals selectively bred for low locomotor responses to novelty exhibited a behavioral phenotype known as “goal-tracking” where they approached the perceived location of a future food reward when a reward-predictive cue was presented. In these animals, dopamine signaling was passive over learning, where no transfer of phasic dopamine signal from US (pellet delivery) to CS (lever extension and light above lever) was observed. By utilizing FLU, the authors found that this subgroup of animals did not display a dopamine dependence in forming a Pavlovian conditioned approach. This effect was in stark contrast to rats bred for high locomotion responses to novelty, who displayed a “sign-tracking” behavioral phenotype where, on cue presentation, animals approached the cue itself rather than the location in which the reward would be delivered. In this group, dopamine signaling was dynamic across learning with phasic dopamine signals transferring from US to CS, and the learning was abolished by dopamine antagonism. Given that senescent animals have drastically reduced dopamine signals without any obvious decline in conditioned approach behavior, we reasoned that senescent animals may have compensated by developing a dopamine independent strategy when learning conditioned approach. In order to test this hypothesis, we employed a similar pharmacological strategy to Flagel, Clark et al (2011). Surprisingly, these experiments revealed that learning of the task was dependent upon dopamine in both the adult and senescent groups. In fact, if anything, the senescent group exhibited greater dependence, as indicated by a greater effect of the lower dose the dopamine receptor antagonist. The greater effect of the low dose of antagonist in the senescent group is likely a direct result of the endogenous dopamine levels being lower. Since FLU is a competitive antagonist, a lower concentration is required for competition when the endogenous ligand is maintained at lower concentrations. To reiterate, the senescent group lost dopamine phasic signaling and yet showed no signs of compensation via a strategy shift (dopamine dependent vs dopamine independent).

How the dopamine-dependent behavior is maintained with such marked decline in phasic dopamine concentration is unclear. It seems that the signaling system may have undergone compensatory changes. It has previously been shown that dopamine stimulated adenylate cyclase activity is not significantly different between adult and senescent animals (Hirschhorn et al, 1982), despite lower receptor levels (Morelli et al., 1990; Suzuki et al, 2001). Another possibility would be that pruning or relocation of receptors has taken place (for conceptual figure see fig. 9). In this scenario dopamine receptors could have moved closer to the synaptic cleft, thus experiencing a higher concentration of volume transmitted dopamine (Garris 1994; Zoli et al., 1999; Cragg and Rice, 2004; Dreyer et al., 2010; Wiencke et al., 2020). However, to date has been no report on whether or not there is dopamine D1 and/or D2 receptor relocalization during aging.

**Figure 9.**
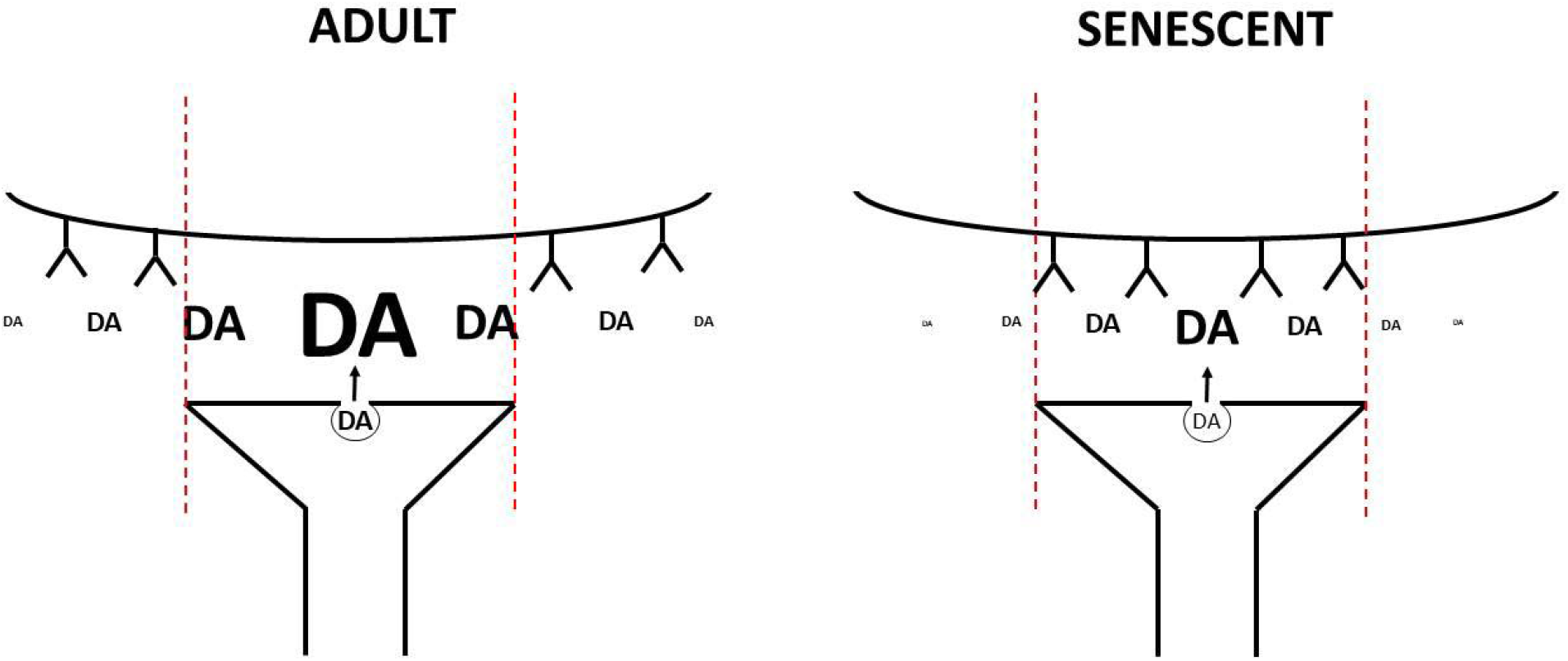
Conceptual drawing of compensatory mechanism underlying observed behavioral responses. **Left panel**, an adult synapse with dopamine receptors extra-synaptically located sensing the dopamine concentration through the diffusion generated concentration gradient. **Right panel**, a senescent synapse with dopamine receptors pruned and concentrated away from extra-synaptic space into synaptic cleft (delineated by red dashed lines). In this way receptors observe similar concentrations despite less being released. This potential relocation into the synaptic cleft could explain the drop in receptor numbers due to the lack of accessibility for anti-bodies.

Overall this work demonstrates that there is an age dependent decrease in phasic dopamine release in male freely behaving F344 rats during a Pavlovian conditioning paradigm. However, no behavioral difference was observed between the two age groups despite marked difference in dopamine transient amplitudes. Subsequent pharmacological manipulation using flupenthixol on a separate cohort revealed that aged animals did not exhibit a shift to a dopamine independent (potentially model-based) learning strategy, but rather displayed increased sensitivity to dopamine antagonism as compared to the adult group. These data indicate that dopamine-dependent learning can be maintained into senescence despite a dramatic reduction in phasic dopamine concentration.

## Acknowledgements

This work was funded by the National Institutes of Health grants R21-AG030775, R01-AG044839, R01-DA027858, and R37-DA051686.

